# COVIDier: A Deep-learning Tool For Coronaviruses Genome And Virulence Proteins Classification

**DOI:** 10.1101/2020.05.03.075549

**Authors:** Peter T. Habib, Alsamman M. Alsamman, Maha Saber-Ayad, Sameh E. Hassanein, Aladdin Hamwieh

## Abstract

COVID-19, caused by SARS-CoV-2 infection, has already reached pandemic proportions in a matter of a few weeks. At the time of writing this manuscript, the unprecedented public health crisis caused more than 2.5 million cases with a mortality range of 5-7%. The SARS-CoV-2, also called novel Coronavirus, is related to both SARS-CoV and bat SARS. Great efforts have been spent to control the pandemic that has become a significant burden on the health systems in a short time. Since the emergence of the crisis, a great number of researchers started to use the AI tools to identify drugs, diagnosing using CT scan images, scanning body temperature, and classifying the severity of the disease. The emergence of variants of the SARS-CoV-2 genome is a challenging problem with expected serious consequences on the management of the disease. Here, we introduce COVIDier, a deep learning-based software that is enabled to classify the different genomes of Alpha coronavirus, Beta coronavirus, MERS, SARS-CoV-1, SARS-CoV-2, and bronchitis-CoV. COVIDier was trained on 1925 genomes, belonging to the three families of SARS retrieved from NCBI Database to propose a new method to train deep learning model trained on genome data using Multi-layer Perceptron Classifier (MLPClassifier), a deep learning algorithm, that could blindly predict the virus family name from the genome of by predicting the statistically similar genome from training data to the given genome. COVIDier able to predict how close the emerging novel genomes of SARS to the known genomes with accuracy 99%. COVIDier can replace tools like BLAST that consume higher CPU and time.

## 1. Introduction

In December 2019, a pneumonia crisis in Wuhan, Hubei Province triggered by an evolved coronavirus emerged and has spread quickly through China at on-going pandemic risk (D. Wang et al. 2020). The pneumonia virus was initially named 2019 novel Coronavirus (2019-nCoV) after virus detection and isolation, which was eventually officially referred to by the WHO as Severe Acute Respiratory Syndrome Coronavirus 2(SARS-CoV-2), (Zhou et al. 2020a). WHO announced a health emergency of international significance on 30 January 2020 with the outbreak of SARS-CoV-2. SARS-CoV-2 has greater transmitting capacity than the SARS-CoV that contributed to an epidemic of SARS in 2003. The exponential growth in reported cases is particularly severe in the detection and treatment in COVID-19. In comparison to certain cases, COVID-19 clinical signs are characterized by respiratory symptoms (Huang et al. 2020). Also, several people with cardiovascular disorders (CVD) are at risk of death (Huang et al. 2020). Respiratory distress syndrome (ARDS) is one of the main causes of respiratory failure and death due to COVID-19 infection (Singhal 2020).

The coronavirus genome ranges between 26,000 and 32,000 bases in size. including from 6 to 11 of open reading frames (ORFs) (Song et al. 2019). The SARS coronaviruses possess four genes that encode the spike(S), the matrix(M), the nucleocapsid (N), and the envelop(E) proteins (Figure 1A).

**Figure 1:**
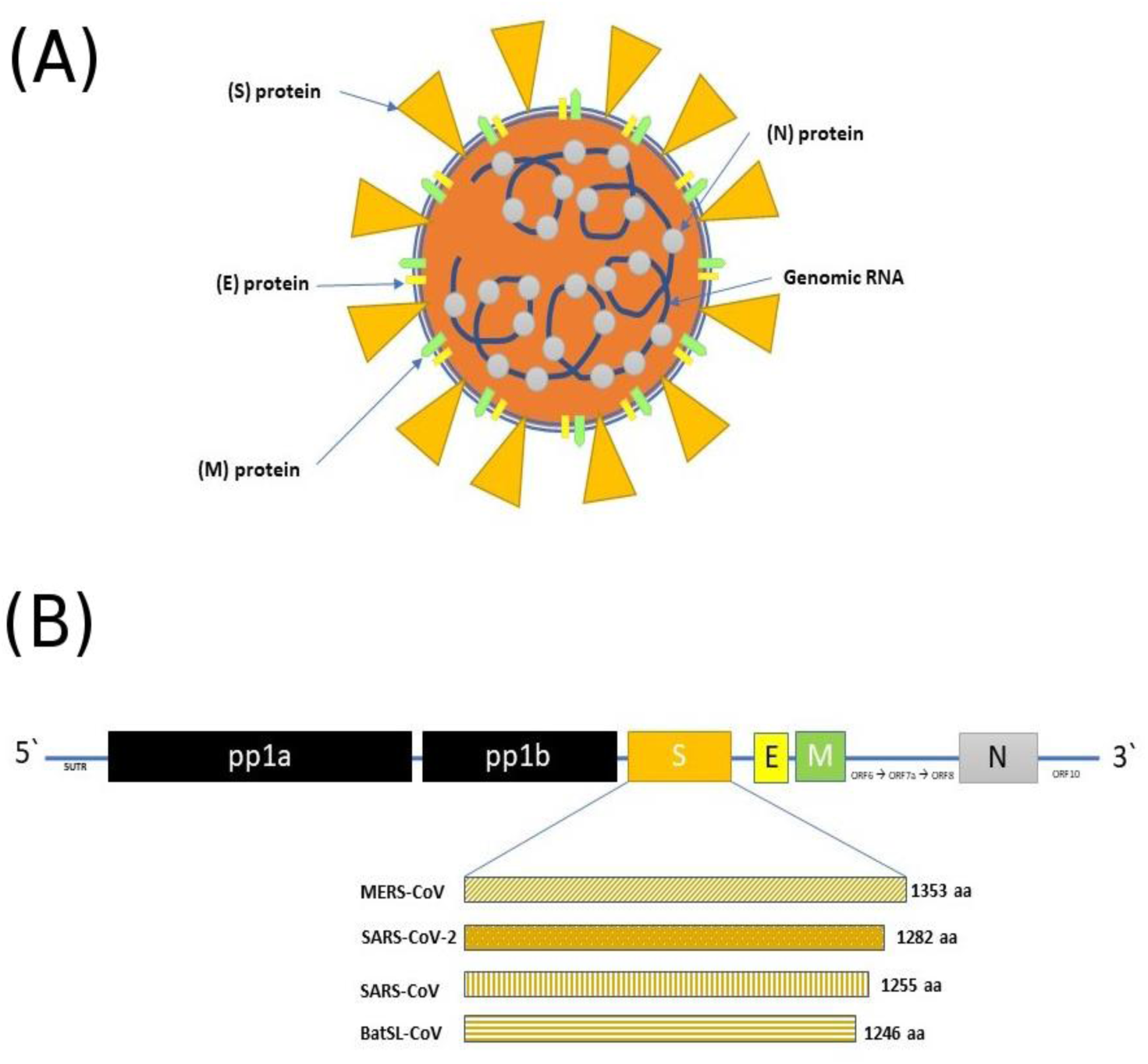
A. Coronavirus structure. B. Genome organization of coronaviruses (subgenus Serbecovirus). The sequence length of each virus Spike (S) protein is shownE=Envelope protein, M= membrane protein, N= Nucleocapsid protein, and S= Spike protein.

The ‘S’ protein is a surface glycoprotein essential for the virus to bind receptors on the host cell, thus determining host tropism (Zhu et al. 2018). Generally, there is low heterogeneity in the genomes of SARS-CoV-2. A hypervariable genomic hotspot is responsible for a Serine/Leucine variation in the viral ORF8-encoded protein (Ceraolo and Giorgi 2020). In the coronavirus genomic analysis, the S gene of SARS-CoV-2 has low sequence identity with other Betacoronaviruses (Figure 1B). The ‘S’ protein in SARS-CoV-2 is longer than SARS-CoV and MERS-CoV. Most importantly are the difference in the receptor-binding domain between the new SARS-CoV-2 and the other two viruses (Zhou et al. 2020b). Different receptors on the host cells are involved in the entry of different viruses, e.g. CD26 fro MERS-CoV and ACE2 for SARS-CoV (Lu et al. 2020). Although several identical residues to that of SARS-CoVs was identified in the SARS-CoV-2, however, they show phylogenic divergence (Letko et al., 2020).

AI and deep learning tools stated to be important for the evaluation and decision-making of patients (Neill 2013). Studies have been conducted to facilitate the diagnosis of disease using AI models (Rajalakshmi et al. 2018; Arabasadi et al. 2017; Kumar et al., 2016; Tomlinson et al. 2009). In the health data collection, the use of smartphones (Ballivian et al. 2015; Braun et al. 2013; Bastawrous and Armstrong 2013; Paolotti et al. 2014) and web-based portals (Fabic et al., 2012) (Metsky et al. 2020) were efficiently examined. These methods will, however, be used in a great time to produce better performance. In addition to the cost-efficiency, the new simulation would help to classify and monitor communities closed to the spreading of the virus. We can also conveniently expand our proposed algorithm to recognize people with moderate symptoms and signs

AI has been reported to be one of the most powerful technologies against the new pandemic. Currently, attempts to create new diagnostic methods using machine learning algorithms have been started. In machine learning, for example, high sensitivity and speed were verified with the SARS-CoV-2 research designs utilizing CRISPR-based viral detection systems (Y. Wang et al. 2020). For screening of COVID-19 patients based on their various breathing patterns, neural network recognition models were developed (Gozes et al. 2020). Likewise, a deep-learning analysis network of the chest CT images was constructed for automatic identification and tracking of COVID-19 patients over time (DeCaprio et al. 2020). Machine learning has been used by DeCapprio et al. to create an initial COVID-19 Vulnerability Index (DeCaprio et al. 2020). The mortality likelihood of a patient developing ARDS is predicted by interrogating the initial symptoms (Munsell et al. 2015).

Machine-learning has previously been used to predict treatment outcomes in various diseases and disorders; for patients with epilepsy (Trebeschi et al. 2019), responses to cancer immunotherapy (Schork 2019). Machine learning now has been used to predict COVID-19 treatment outcomes (Broecker and Moelling 2019). Viruses help human immune systems evolve over long decades. An image emerges from and is transported by viruses and transposable elements into the various cellular immune systems. Immune systems have possibly evolved from simple exclusion from superinfection to very complex defense strategies (CDCP 2017). However, the concept of ‘viral genome - host Genome interaction’ continues to be the trend after reporting many cellular infection processes are mediated by the virus and the host through a network of genes. The interaction? between genome and genome naturally influences the immune response as well as the drug response. For instance, infection with some influenza A virus subtypes such as H5N1 and H7N9 may cause severe respiratory disease (Berhane et al. 2016; Josset et al. 2014). Previous studies showed that weaker innate immune responses were induced by H5N1 and H7N9 (Samy et al. 2016; Suzuki et al. 2009). At the drug response stage, drug sensitivity testing demonstrated that the new isolates were oseltamivir-resistant but peramivir-sensitive (Gao et al. 2014).

Classifying the genomes according to their immune and drug response will become increasingly important for future personalization of the diagnosis and treatment by viral genome sort. Here we introduce COVIDier, a deep learning tool to classify SARS subtypes genomes to the main families: Alpha coronavirus, Beta coronavirus, MERS, SARS-CoV-1, SARS-CoV-2, and bronchitis-CoV. COVIDier is a user-friendly and open-source software that can be modified and integrated into different frameworks.

## 2. Implementation

COVIDier is written in Python, one of the most used programming languages in the bioinformatics that is easy to learn, powerful, and has tremendous useful libraries including Biopython, Pandas and ETE3. COVIDier was implemented as a Python tool with open source code to be modified and integrated into a different frame.

### 2.1. Data Collection

Training and testing data sets include a 1925 genome of Alpha coronavirus, Beta coronavirus, MERS, SARS-CoV-1, SARS-CoV-2, and bronchitis-CoV and 7702 protein of spike protein, envelope protein, nucleocapsid phosphoprotein, membrane glycoprotein, a surface glycoprotein from the previously mentioned genomes. distributed data for different coronaviridae were collected from the NCBI. The causative agents of SARS and COVID-19, and the associated family that causes bat SARS, are especially of interest. A CSV file of genomic sequence with NCBI accession number contains the data unique to those three classes.

### 2.2. Data pre-processing

We used the CounterVectorizer function to convert a set of text to a token count matrix, as our data was expressed in text form. We used CounterVectorizer to convert textual data to numeric in genome sequence to allow the algorithm to learn and improve our deep learning model’s predictive potential. To vectorize the data, we deploy the training file as control or/and normal.

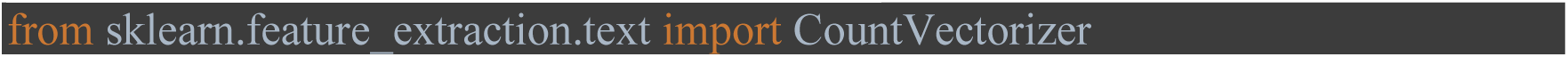

### 2.3. Algorithm Selection

We used the selection code from the TarDict algorithm to find the most suitable algorithm on the genome data (Habib et al. 2020). By looping 23 algorithms. Testing and evaluation showed that MLPClassifier is the best 99.6% accuracy.

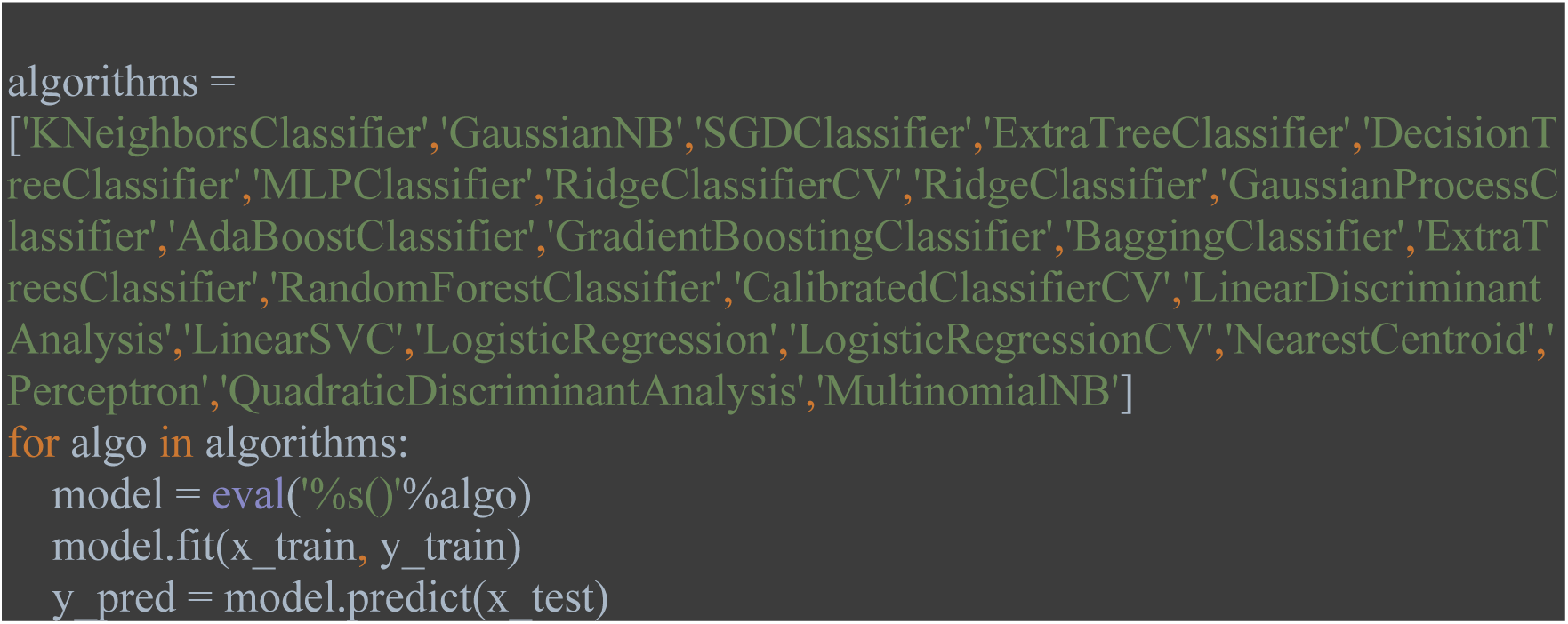

### 2.4. Multi-layer Perceptron Classifier (MLPClassifier)

MLPClassifier can be presented as generalizations of linear models executing multiple processing stages to arrive at a decision. Linear regression defined as:

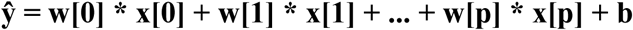

Where ŷ is the sum of the input features (x[0]) to (x[p]), weighted by the learned coefficients (w[0]) to (w[p]). MLPClassifier trains using Back-propagation introduced by Rumelhart et., al in 1989. The algorithm is used to effectively train a neural network through a method called chain rule. While training, after each forward pass through a network, backpropagation performs a backward pass while adjusting the model’s parameters (weights and biases).

The 8-layer neural network consists of 1925 neurons for input layer, 6 hidden layers each with 150,250,260,250,180,70 neurons respectively, and 1neuron of output layer for the input layer, 4 neurons for the hidden layers and 1 neuron for the output layer as shown in **Figure 2**.

**Figure 2:**
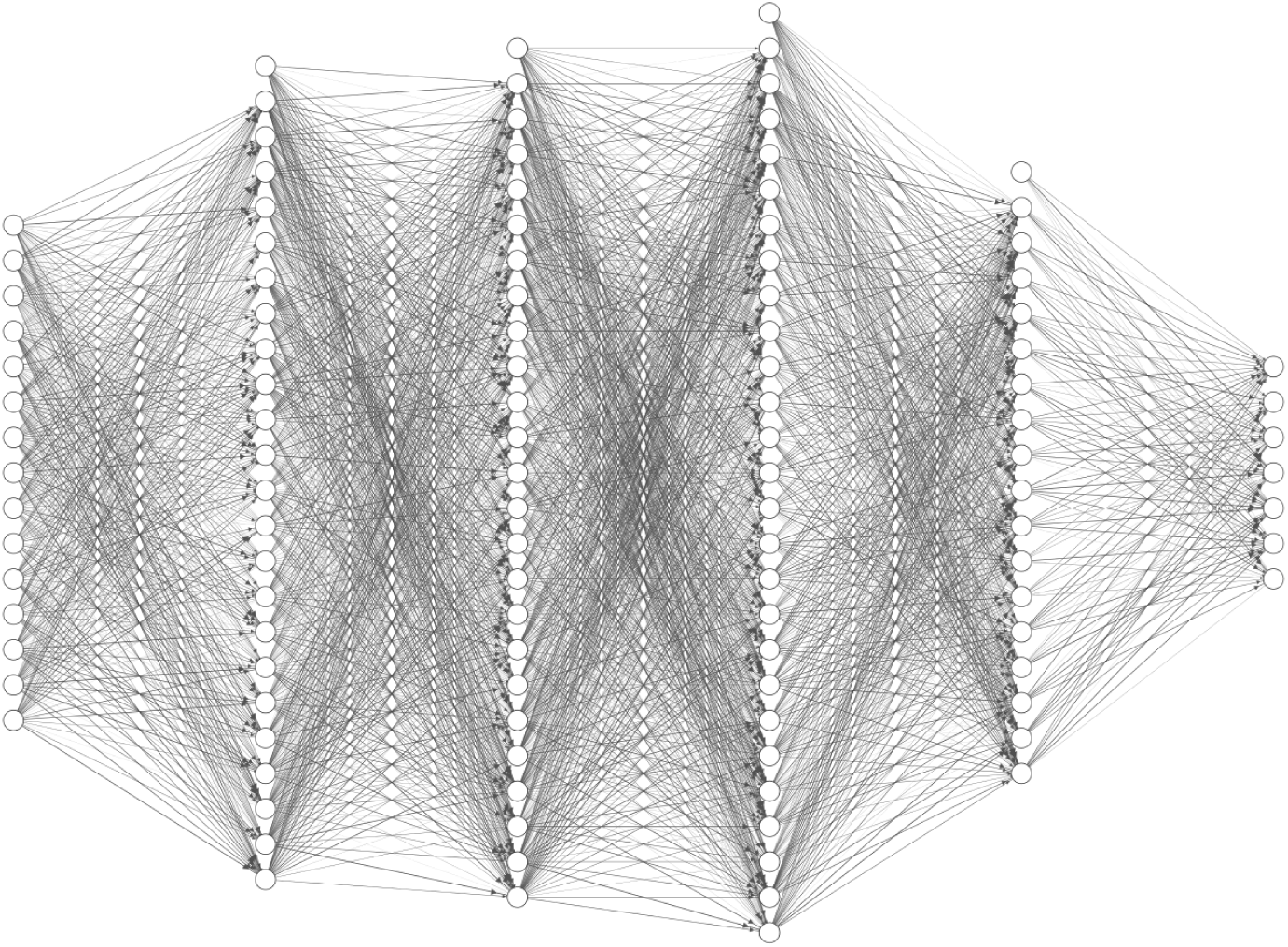
The architecture of the neural network (number of neurons in figure divided over 100 to be represent-able).

For each forward pass, the final step is to compare the estimated output x against the predicted output y. The output y is part of the training dataset (x, y) where x is the origin. Evaluation between x and y occurs by a cost function. This can be as basic as mean squared error (MSE) or more complex as cross-entropy. The model was designed to know how much to change its parameters to get closer to the expected output y, based on cost-function meaning. This happens using the Backpropagation algorithm.

Backpropagation aims to reduce the cost function by adjusting the network’s weights and biases. The degree of adjustment is determined by the cost function concerning certain parameters. Firstly, The algorithm select values of w and b are random, the initial learning rate of Epsilon (e) which is 0.001 in our model defines the effect of the gradient. The weight or bias derivative of the cost function can be determined using partial cost function derivatives in the individual weights or biases. Learning terminates until the cost function is reduced.

### 2.5. Functions

COVIDier consisting of 5 modules to receive user input whether in FASTA format or list of accession numbers, predict the family of the genome, perform multiple sequence alignment, and visualize the alignment.

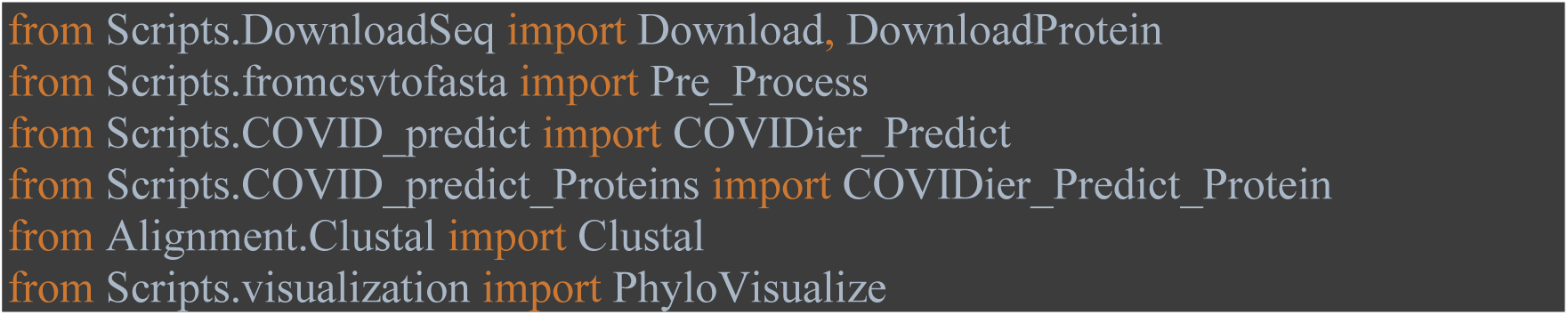

Download() for nucleotide and DownloadProtein() for protein are functions imported from the Biopython library to download the record of given IDs in the list. Pre_Process() is a regex-based function to extract IDs and sequences from the FASTA file to prepare it in the proper format of prediction. COVIDier_Predict() function that reads the given nucleotide genomes, vectorizes, and predicts the family of the genome and COVID_Predict_Protein() do the same but for protein. Clustal() is a part from the Biopython library for multiple sequence alignment used in COVIDier to show the differences between the input genomes. PhyloVisualize() uses the ETE3 python package to display the multiple sequence alignment produced by Clustal().

## 3. Results and Discussion

The COVIDier is a deep learning tool that uses Sci-kit learn packages (Pedregosa et al. 2011) to classify between Alpha coronavirus, Beta coronavirus, MERS, SARS-CoV-1, SARS-CoV-2, and bronchitis-CoV genomes with accuracy 99.6% and visualize the differences between input genomes by browsing the multiple sequence alignment. Moreover, COVIDier provides users with genome length and count each nucleotide content in the genome. It consists of 8 layers, 1 for input, 1 for output, and 6 hidden layers as shown in **Figure 2**.

### 3.1. Confusion Matrix and Classification Report

In the field of machine learning, and in particular the problem of statistical classification, a confusion matrix, also known as an error matrix, is a description of predictive effects on a classification problem. The number of observations that are accurate and inaccurate is listed with count values and broken down by class. The confusion matrix reveals how the classification model becomes confused when making predictions. This provides insight not only into the errors produced by a classifier but also, more critically, into the types of errors produced. The accuracy of predictions from a classification algorithm is evaluated using a Classification Report. How many are True Predictions and how many are Wrong. True Positives, False Positives, True negatives, and False Negatives are used to estimate a classification report’s metrics as shown in **Tables 1** and **2**.

**Table 1:** Classification report indicates each class of SARS genomes. Precision is a classifier’s capacity not to mark a positive instance which is in fact negative. Recall is a classifier’s capacity to identify all appropriate cases. The F1 value is a weighted, harmonic accuracy and recall measure.

| Virus genome | precision | recall | f1-score |
| --- | --- | --- | --- |
| SARS-CoV-1 | 1.00 | 0.99 | 0.99 |
| SARS-CoV-2 | 1.00 | 1.00 | 1.00 |
| Alpha-Cov | 0.99 | 1.00 | 0.99 |
| Beta-Cov | 0.98 | 0.99 | 0.99 |
| MERS | 0.99 | 1.00 | 0.99 |
| Bronchitis | 1.00 | 0.99 | 0.99 |
| accuracy |  |  | 0.99 |
| macro avg | 0.98 | 0.98 | 0.99 |
| weighted avg | 0.99 | 0.99 | 0.99 |

**Table 2:** Classification report indicates each class of SARS proteins. Precision is a classifier’s capacity not to mark a positive instance which is, in fact, negative. The recall is a classifier’s capacity to identify all appropriate cases. The F1 value is a weighted, harmonic accuracy, and recalls measure.

| Virus protein | Precision | Recall | f1-score |
| --- | --- | --- | --- |
| Spike protein | 0.96 | 0.99 | 0.99 |
| Nucleocapsid Phosphoprotein | 1 | 1 | 1 |
| Envelope Protein | 1 | 0.99 | 0.99 |
| Membrane Glycoprotein | 0.99 | 1 | 0.99 |
| Surface Glycoprotein | 0.98 | 1 | 0.98 |
| accuracy |  |  | 0.98 |
| macro avg | 0.97 | 0.98 | 0.99 |
| weighted avg | 0.98 | 0.99 | 0.99 |

### 3.2. Use case 1: Classification of SARS Genomes or list of IDs

The following command line can be used to identify a large number of SARS genomes and visualize the alignment or IDs list of protein (spike protein in this case) and visualize the alignment shown in **Figures 3** and **4**.

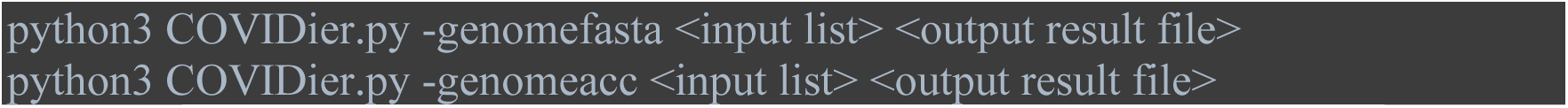

**Figure 3:**
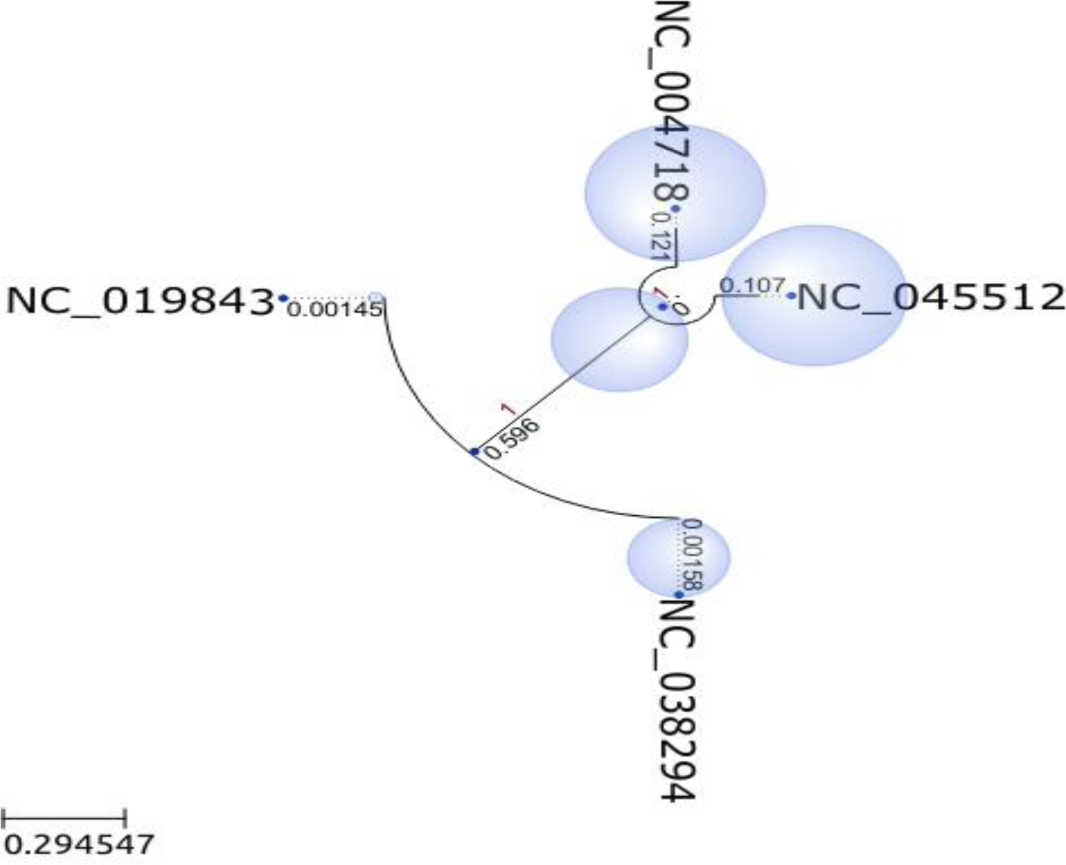
Bubble tree maps of different genomes

**Figure 4:**
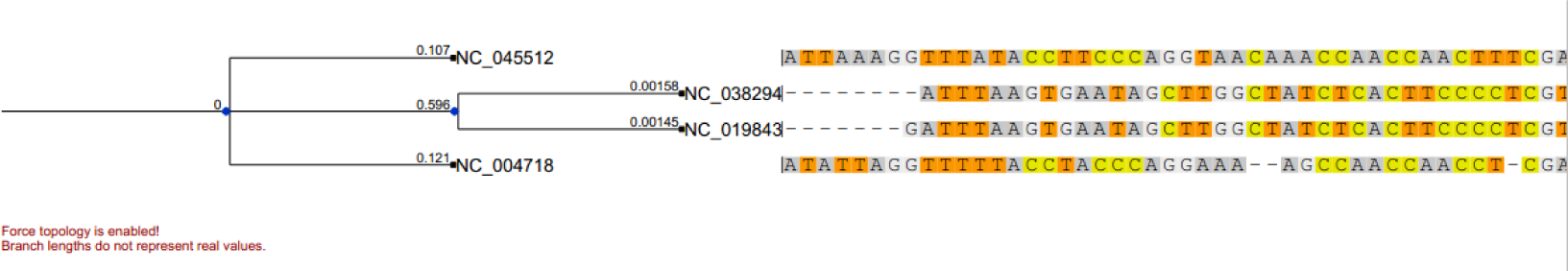
PhyloTree with in-place alignment and evolutionary events of different genomes.

### 3.3. Use case 2: Classification of SARS protein or the list of IDs

The following two command lines can be used to identify a large number of unknown genomes or IDs of known protein (spike protein in this case) and visualize the alignment shown in **Figures** 5 and **6**.

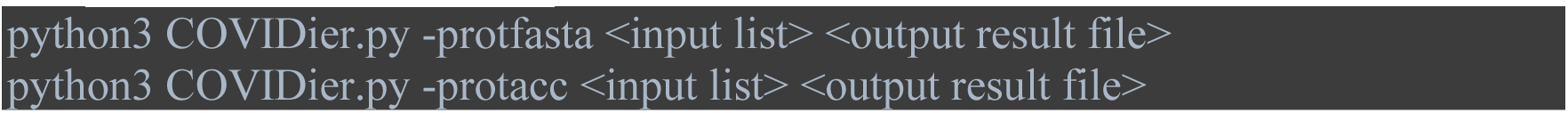

**Figure 5:**
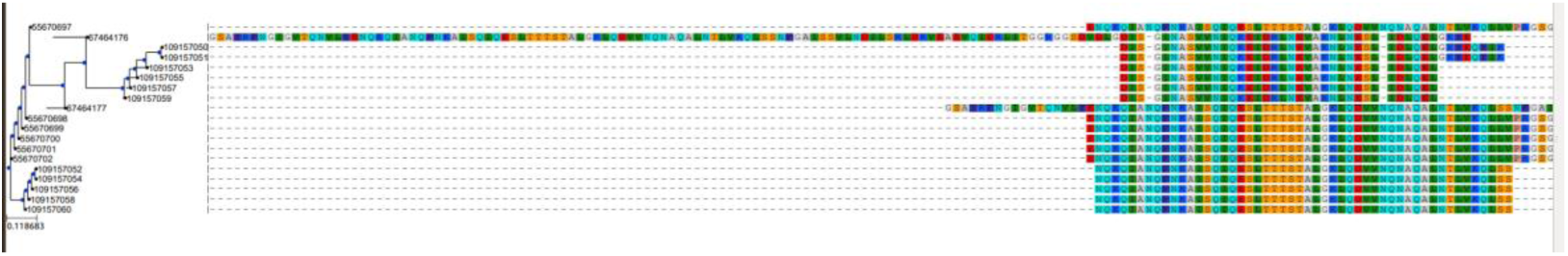
PhyloTree with in-place alignment and evolutionary events of proteins genomes.

**Figure 6:**
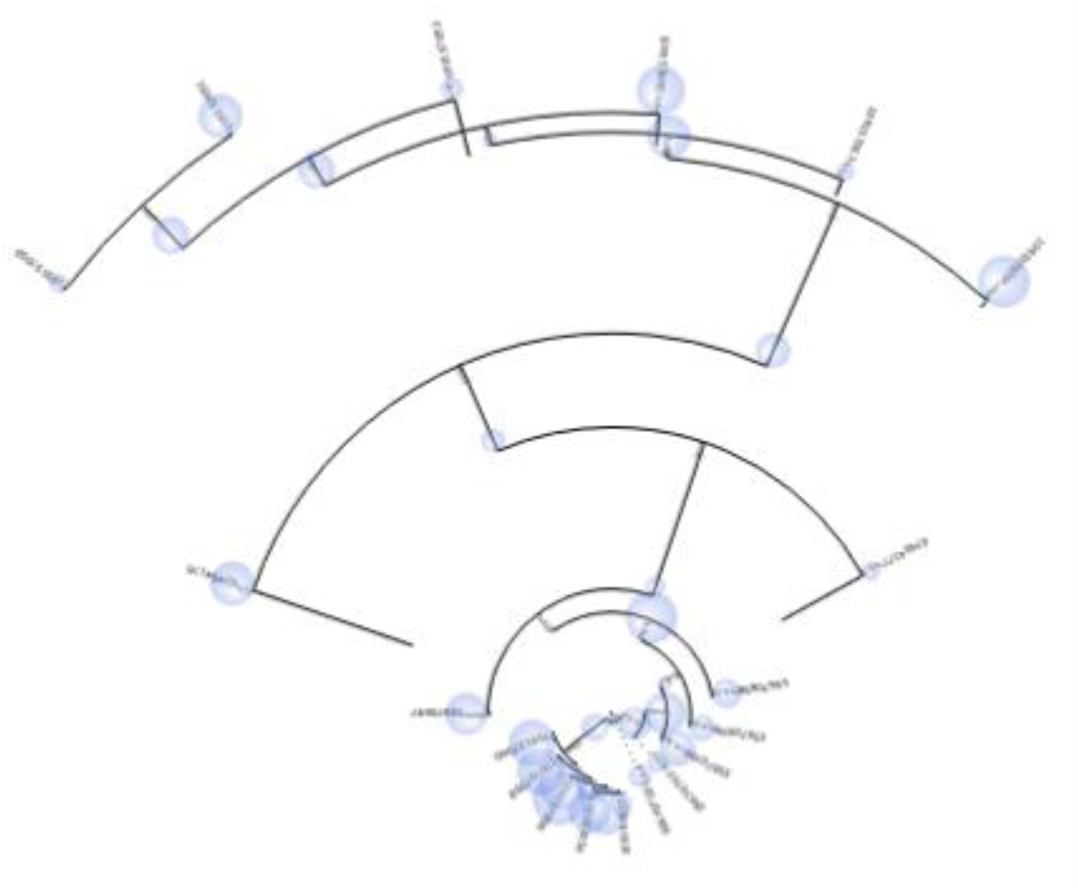
Bubble treemaps of proteins genomes

### 3.2. COVIDier and BLAST

A well-known tool that widely used for alignment and classification of genomes is BLAST (Altschul et al. 1990). We used the 450 of training genomes to test the performance of COVIDier and BLAST. The results show that COVIDier has much higher performance and speed where it was able to classify 450 genomes in 18 seconds, while BLAST continues the alignment for 5 minutes and collapsed. Such results may raise the importance of using such an approach instead of BLAST for genomic classification on a large scale.

## 4. State-of-The-Art

From the beginning of COVID-19, authorities over the world spent their efforts on fighting the virus, or at least slow its spreading. Researchers are called to accelerate the pace of their efforts using AI to stop the pandemic. COVIDier deploying deep learning able to classify different genomes of SARS in fast and accurate manner focusing the light on the importance of AI in such processes the require high computation power and the need to replace the traditional tools with artificially intelligent tools to maximize the accuracy and minimize the required time and processing power. Detecting the variants in the virus genome with high accuracy may lead to better population-based vaccine designing (according to the geographical location), and hopefully to better prevention of the disease.

## 5. Conclusion

There has been a sparkling spread of the atypical viral pneumonia caused by the zoonotic coronavirus SARS-CoV-2. The pandemic is unprecedented for health care systems worldwide. Detection of the specific genome sequence of the virus is highly important. A deep learning model was trained on genome data using Multi-layer Perceptron Classifier (MLPClassifier), a deep learning algorithm, that could blindly predict the virus family name from the genome through statistically similar genome from training data to the given genome. Accordingly, we designed “COVIDier”, a CPU- and time-saving tool, to predict variability in coronavirus genome with high accuracy. In this paper, we used 1925 genomes belong to the three families of SARS retrieved from NCBI Database to propose a new method to train deep learning model trained on genome data using Multi-layer Perceptron Classifier (MLPClassifier), a deep learning algorithm, that could blindly predict the virus family name from the genome of by predicting the statistically similar genome from training data to the given genome. A tool such as COVIDier predicted to replace tools like BLAST that consume higher CPU and time.

## 6. Data Availability

Source code freely available on Github: https://github.com/peterhabib/COVIDier

## References

Altschul, Stephen F, Warren Gish, Webb Miller, Eugene W Myers, and David J Lipman. 1990. “Basic Local Alignment Search Tool.” Journal of Molecular Biology 215 (3): 403–10.

Arabasadi, Zeinab, Roohallah Alizadehsani, Mohamad Roshanzamir, Hossein Moosaei, and Ali Asghar Yarifard. 2017. “Computer Aided Decision Making for Heart Disease Detection Using Hybrid Neural Network-Genetic Algorithm.” Computer Methods and Programs in Biomedicine 141: 19–26.

Ballivian, Amparo, J Azevedo, Will Durbin, J Rios, J Godoy, and C Borisova. 2015. “Using Mobile Phones for High-Frequency Data Collection.” Mobile Research Methods, 21.

Bastawrous, Andrew, and Matthew J Armstrong. 2013. “Mobile Health Use in Low- and High-Income Countries: An Overview of the Peer-Reviewed Literature.” Journal of the Royal Society of Medicine 106 (4): 130–42.

Berhane, Yohannes, Darwyn Kobasa, Carissa Embury-Hyatt, Brad Pickering, Shawn Babiuk, Tomy Joseph, Victoria Bowes, et al. 2016. “Pathobiological Characterization of a Novel Reassortant Highly Pathogenic H5N1 Virus Isolated in British Columbia, Canada, 2015.” Scientific Reports 6 (1): 1–15.

Braun, Rebecca, Caricia Catalani, Julian Wimbush, and Dennis Israelski. 2013. “Community Health Workers and Mobile Technology: A Systematic Review of the Literature.” PloS One 8 (6).

Broecker, Felix, and Karin Moelling. 2019. “Evolution of Immune Systems from Viruses and Transposable Elements.” Frontiers in Microbiology 10: 51.

Ceraolo, Carmine, and Federico M Giorgi. 2020. “Genomic Variance of the 2019-NCoV Coronavirus.” Journal of Medical Virology.

DeCaprio, Dave, Joseph Gartner, Thadeus Burgess, Sarthak Kothari, and Shaayan Sayed. 2020. “Building a COVID-19 Vulnerability Index.” ArXiv Preprint ArXiv:2003.07347.

Fabic, Madeleine Short, YoonJoung Choi, and Sandra Bird. 2012. “A Systematic Review of Demographic and Health Surveys: Data Availability and Utilization for Research.” Bulletin of the World Health Organization 90: 604–12.

for Disease Control, Centers, Prevention, and others. 2017. “Avian Influenza A (H7N9) Virus| Avian Influenza (Flu)[WWW Document].” URL https://www.Cdc.Gov/Flu/Avianflu/H7n9-Virus.Htm (Accessed 2.9. 17).

Gao, Hai Nv, Hang Ping Yao, Wei Feng Liang, Xiao Xin Wu, Hai Bo Wu, Nan Ping Wu, Shi Gui Yang, et al. 2014. “Viral Genome and Antiviral Drug Sensitivity Analysis of Two Patients from a Family Cluster Caused by the Influenza A (H7N9) Virus in Zhejiang, China, 2013.” International Journal of Infectious Diseases 29: 254–58.

Gozes, Ophir, Maayan Frid-Adar, Hayit Greenspan, Patrick D Browning, Huangqi Zhang, Wenbin Ji, Adam Bernheim, and Eliot Siegel. 2020. “Rapid Ai Development Cycle for the Coronavirus (Covid-19) Pandemic: Initial Results for Automated Detection & Patient Monitoring Using Deep Learning Ct Image Analysis.” ArXiv Preprint ArXiv:2003.05037.

Habib, Peter T, Alsamman M Alsamman, Sameh E Hassanein, and Aladdin Hamwieh. 2020. “TarDict: RandomForestClassifier-Based Software Predict Drug-Tartget Interaction.” BioRxiv.

Huang, Chaolin, Yeming Wang, Xingwang Li, Lili Ren, Jianping Zhao, Yi Hu, Li Zhang, et al. 2020. “Clinical Features of Patients Infected with 2019 Novel Coronavirus in Wuhan, China.” The Lancet 395 (10223): 497–506.

Josset, Laurence, Hui Zeng, Sara M Kelly, Terrence M Tumpey, and Michael G Katze. 2014. “Transcriptomic Characterization of the Novel Avian-Origin Influenza A (H7N9) Virus: Specific Host Response and Responses Intermediate between Avian (H5N1 and H7N7) and Human (H3N2) Viruses and Implications for Treatment Options.” MBio 5 (1): e01102--13.

Kumar, Vinayshekhar Bannihatti, Sujay S Kumar, and Varun Saboo. 2016. “Dermatological Disease Detection Using Image Processing and Machine Learning.” In 2016 Third International Conference on Artificial Intelligence and Pattern Recognition (AIPR), 1–6.

Letko, Michael, Andrea Marzi, and Vincent Munster. 2020. “Functional Assessment of Cell Entry and Receptor Usage for SARS-CoV-2 and Other Lineage B Betacoronaviruses.” Nature Microbiology 5 (4): 562–69.

Lu, Roujian, Xiang Zhao, Juan Li, Peihua Niu, Bo Yang, Honglong Wu, Wenling Wang, et al. 2020. “Genomic Characterisation and Epidemiology of 2019 Novel Coronavirus: Implications for Virus Origins and Receptor Binding.” The Lancet 395 (10224): 565–74.

Metsky, Hayden C, Catherine A Freije, Tinna-Solveig F Kosoko-Thoroddsen, Pardis C Sabeti, and Cameron Myhrvold. 2020. “CRISPR-Based COVID-19 Surveillance Using a Genomically-Comprehensive Machine Learning Approach.” BioRxiv.

Munsell, Brent C, Chong-Yaw Wee, Simon S Keller, Bernd Weber, Christian Elger, Laura Angelica Tomaz da Silva, Travis Nesland, Martin Styner, Dinggang Shen, and Leonardo Bonilha. 2015. “Evaluation of Machine Learning Algorithms for Treatment Outcome Prediction in Patients with Epilepsy Based on Structural Connectome Data.” Neuroimage 118: 219–30.

Neill, Daniel B. 2013. “Using Artificial Intelligence to Improve Hospital Inpatient Care.” IEEE Intelligent Systems 28 (2): 92–95.

Paolotti, Daniela, Annasara Carnahan, Vittoria Colizza, Ken Eames, J Edmunds, Gabriel Gomes, C Koppeschaar, et al. 2014. “Web-Based Participatory Surveillance of Infectious Diseases: The Influenzanet Participatory Surveillance Experience.” Clinical Microbiology and Infection 20 (1): 17–21.

Pedregosa, Fabian, Gaël Varoquaux, Alexandre Gramfort, Vincent Michel, Bertrand Thirion, Olivier Grisel, Mathieu Blondel, et al. 2011. “Scikit-Learn: Machine Learning in Python.” Journal of Machine Learning Research 12 (Oct): 2825–30.

Rajalakshmi, Ramachandran, Radhakrishnan Subashini, Ranjit Mohan Anjana, and Viswanathan Mohan. 2018. “Automated Diabetic Retinopathy Detection in Smartphone-Based Fundus Photography Using Artificial Intelligence.” Eye 32 (6): 1138–44.

Samy, Ahmed A, Mona I El-Enbaawy, Ahmed A El-Sanousi, Soad A Nasef, Mahmoud M Naguib, E M Abdelwhab, Hirokazu Hikono, and Takehiko Saito. 2016. “Different Counteracting Host Immune Responses to Clade 2.2. 1.1 and 2.2. 1.2 Egyptian H5N1 Highly Pathogenic Avian Influenza Viruses in Naive and Vaccinated Chickens.” Veterinary Microbiology 183: 103–9.

Schork, Nicholas J. 2019. “Artificial Intelligence and Personalized Medicine.” in Precision Medicine in Cancer Therapy, 265–83. Springer.

Singhal, Tanu. 2020. “A Review of Coronavirus Disease-2019 (COVID-19).” The Indian Journal of Pediatrics, 1–6.

Song, Zhiqi, Yanfeng Xu, Linlin Bao, Ling Zhang, Pin Yu, Yajin Qu, Hua Zhu, Wenjie Zhao, Yunlin Han, and Chuan Qin. 2019. “From SARS to MERS, Thrusting Coronaviruses into the Spotlight.” Viruses 11 (1): 59.

Suzuki, Koutaro, Hironao Okada, Toshihiro Itoh, Tatsuya Tada, Masaji Mase, Kikuyasu Nakamura, Masanori Kubo, and Kenji Tsukamoto. 2009. “Association of Increased Pathogenicity of Asian H5N1 Highly Pathogenic Avian Influenza Viruses in Chickens with Highly Efficient Viral Replication Accompanied by Early Destruction of Innate Immune Responses.” Journal of Virology 83 (15): 7475–86.

Tomlinson, Mark, Wesley Solomon, Yages Singh, Tanya Doherty, Mickey Chopra, Petrida Ijumba, Alexander C Tsai, and Debra Jackson. 2009. “The Use of Mobile Phones as a Data Collection Tool: A Report from a Household Survey in South Africa.” BMC Medical Informatics and Decision Making 9 (1): 51.

Trebeschi, S, S G Drago, N J Birkbak, I Kurilova, A M Calin, A Delli Pizzi, F Lalezari, et al. 2019. “Predicting Response to Cancer Immunotherapy Using Noninvasive Radiomic Biomarkers.” Annals of Oncology 30 (6): 998–1004.

Wang, Dawei, Bo Hu, Chang Hu, Fangfang Zhu, Xing Liu, Jing Zhang, Binbin Wang, et al. 2020. “Clinical Characteristics of 138 Hospitalized Patients with 2019 Novel Coronavirus--Infected Pneumonia in Wuhan, China.” Jama 323 (11): 1061–69.

Wang, Yunlu, Menghan Hu, Qingli Li, Xiao-Ping Zhang, Guangtao Zhai, and Nan Yao. 2020. “Abnormal Respiratory Patterns Classifier May Contribute to Large-Scale Screening of People Infected with COVID-19 in an Accurate and Unobtrusive Manner.” ArXiv Preprint ArXiv:2002.05534.

Zhou, Peng, Xing-Lou Yang, Xian-Guang Wang, Ben Hu, Lei Zhang, Wei Zhang, Hao-Rui Si, et al. 2020a. “A Pneumonia Outbreak Associated with a New Coronavirus of Probable Bat Origin.” Nature 579 (7798): 270–73.

Zhou, Peng. 2020b. “Discovery of a Novel Coronavirus Associated with the Recent Pneumonia Outbreak in Humans and Its Potential Bat Origin.” BioRxiv.

Zhu, Zhaozhong, Zheng Zhang, Wenjun Chen, Zena Cai, Xingyi Ge, Haizhen Zhu, Taijiao Jiang, Wenjie Tan, and Yousong Peng. 2018. “Predicting the Receptor-Binding Domain Usage of the Coronavirus Based on Kmer Frequency on Spike Protein.” Infection, Genetics and Evolution 61: 183.

